# Size Structures Sensory Hierarchy in Ocean Life

**DOI:** 10.1101/018937

**Authors:** Erik A. Martens, Navish Wadhwa, Nis S. Jacobsen, Christian Lindemann, Ken H. Andersen, André Visser

## Abstract

Survival in aquatic environments requires organisms to have effective means of collecting information from their surroundings through various sensing strategies. In this study, we explore how sensing mode and range depend on body size. We find a hierarchy of sensing modes determined by body size. With increasing body size, a larger battery of modes becomes available (chemosensing, mechanosensing, vision, hearing, and echolocation, in that order) while the sensing range also increases. This size-dependent hierarchy and the transitions between primary sensory modes are explained on the grounds of limiting factors set by physiology and the physical laws governing signal generation, transmission and reception. We theoretically predict the body size limits for various sensory modes, which align well with size ranges found in literature. The treatise of all ocean life, from unicellular organisms to whales, demonstrates how body size determines available sensing modes, and thereby acts as a major structuring factor of aquatic life.

## I. INTRODUCTION

The marine pelagic environment is sparsely populated. To survive, organisms must scan volumes of water millions of times their own body volumes per day [1]. While searching is a challenge in itself, there is also the continual risk of predation. The result is a strong evolutionary drive to effectively gather information on the proximity of prey, mates and predators [2]. Here, we examine the means by which this information is gathered by marine pelagic organisms, that is, their sensory ability. In particular, we wish to understand relationships between the size of an organism and the usability of the various types of senses.

Indeed, size is a key parameter to characterize biological processes in marine environments [1, 3–6]. A cursory examination indicates at least some size-dependent organization as to which sensory modes organisms use in the marine pelagic environment. For instance, the smallest organisms (e.g., bacteria) depend heavily on chemical signals, while for larger animals (e.g., copepods), sensing of fluid flows becomes important, too. For even larger organisms, vision (e.g., crustaceans and fish), hearing (e.g., fish) and echolocation (e.g., toothed whales) become increasingly relevant sensory modes (Supplementary Figure 1). How can we understand this pattern on the grounds of physiology and physics using scaling rules, which are the two basic constraints on the workings of any organism [7, 8]? Our aim here is to determine the body size limits of different sensing modes based on physical grounds, and to explain how the sensory hierarchy is structured by size.

## II. SENSING AS A PHYSICAL PROCESS

Our goal is to understand how size determines sensory modes available to an organism. We restrict ourselves to those sensory modes that are the primary means of remotely detecting the presence of other organisms: chemosensing of compounds, mechanosensing of flow disturbances provoked by moving animals, image vision in sufficiently lit areas, hearing of sound waves, and their generation for echolocation. We further restrict ourselves to the pelagic zone. All sensing involves an organism and a target; thus, we refer to the *organism* of size *L* and the *target* of size *L*_t_. The two lengths are related via the dimensionless size preference *p* = *L*_t_*/L* (we assume *p* = 0.1 for predation, *p* = 1 for mating, *p* = 10 for predator avoidance). Clearly, other modes such as electroreception [9] or magnetoreception [10] may supplement the above mentioned modes, and organisms may switch between sensing modes depending on proximity to the target; here, however, we restrict ourselves to the aforementioned senses and consider them as the predominant primary sensory modes.

It is possible to decompose sensing into three fundamental sub-processes (Figure 1):

### Generation

Animals emit signals by creating fluid disturbances, creating sounds or reflecting ambient light. The target’s features such as its size, *L*_t_, affect the signal. Chemosensing, hearing and mechanosensing require a signal or an action from the target, whereas vision and echolocation do not. Echolocation in particular is an ‘active sense’, as the signal is generated by the organism and hence influenced by organism features such as size *L*.

### Propagation

The distance over which a signal propagates before getting subdued by noise is sensitive to many factors. For instance, the oceans are awash with traces of various chemicals. Detection of a specific compound requires concentrations higher than the background, and depends on its diffusivity, release rate, stability, etc. This distance sets a sensing range *R*.

### Detection

Is the organism — given the physical constraints — able to build a sensor? This requires a costeffective mechanism by which information can be collected at a practical level of resolution. Size and complexity of the organism determine this ability.

**FIG. 1.**
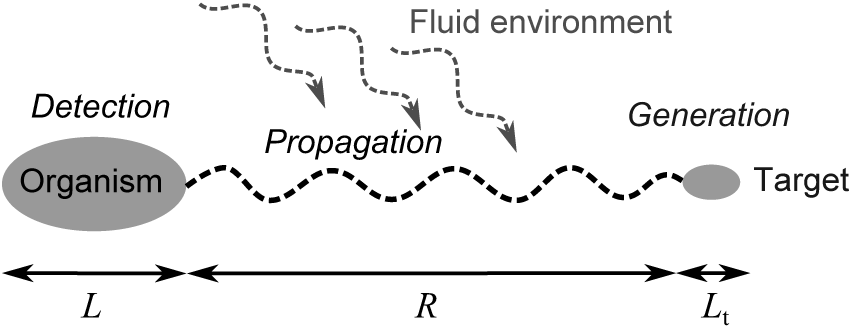
Schematic of the participants and the processes involved in sensing.

Each of these sub-processes is constrained by size. Thus the length scale imprints itself automatically on the remote detection of other organisms. But limits of the usage of specific sensing modes are not necessarily clear-cut. For instance, in case of vision, the boundary between an image-forming eye (e.g., in fish) and nonimage forming ‘eye spots’ that enable phototaxis (e.g., in copepods, protists) is not sharply defined. Moreover, simultaneous use of multiple senses complicates the situation. We make the simplifying assumption of no integration between senses, and treat them in isolation from each other. Within its limitations, this investigation may not yield exact numbers; it provides *characteristic* body-size limits for the sensory modes and yields valuable understanding of the structure of sensing in marine life, based on first principles.

## III. CHEMOSENSING

The ability to detect chemical compounds is ubiquitous. All life forms have this ability and are equipped with chemosensing apparatuses [11]. Chemotaxis and the use of chemosensing in remote detection can be divided into two modes: i) gradient climbing defined as moving along a gradient towards (or away from) a stationary target, and ii) following a trail laid out by a moving target [12, 13].

### A. Size limits for chemosensing

Gradient climbing ability would be size independent, were it not for two randomizing physical effects. For very small organisms, gradient climbing ability is impaired due to Brownian rotation [14], caused by molecular motions in the fluid. Due to this, the organism cannot direct itself along a gradient using a biased random walk (Figure 2A). This happens for *L* less than the length scale characteristic of Brownian motion, *L*_Br_ (0.1 *−*1*μ*m) [15]. Using a similar argument, Dusenbery [16] has argued that below *L* = 0.6 *μ*m, directed motility, and thus chemotaxis, is infeasible due to Brownian rotation.

An upper limit for gradient climbing is imposed when turbulence disrupts the smoothness of the chemical gradient, for *L* greater than the Batchelor scale *L*_B_ *≈*(*vD*^2^/ε)^1/4^, where *v* is the kinematic viscosity, *D* the molecular diffusivity, and ε the turbulent energy dissipation rate. *L*_B_ is the length scale at which the diffusion time scale becomes comparable to the dissipation time for the smallest turbulent eddies (Figure 2B). In the ocean, ε ranges between 10^−8^ and 10^−3^ m^2^s^−3^ [17, 18]. *L*_B_ is between 5 and 100 *μ*m in moderate turbulence (for a typical value of *D ∼* 10^−9^ m^2^s^−1^), but can become much larger in quiescent environments.

For detecting a moving target that releases a chemical trail, the physical constraints are similar to gradient climbing. For *L* above the Kolmogorov scale *L*_K_ *≈*(*v*^3^/ε)^1/4^, directional information in the trail is reduced due to the isotropy in turbulent flows [19], impairing chemotaxis. *L*_K_ is around 1 cm in moderate turbulence [17], above which trail following becomes progressively worse. When *L* is larger than the integral length scale *L*_I_, trail following may become effective again as the turbulent trail at this scale is anisotropic (Figure 2C). Typical values for *L*_I_ in a stratified ocean are around 1 m or larger [20, 21]. Thus, between *∼* 1 cm and *∼*1 m, trail following is impaired, and requires averaging over space and time [22]. Note that in the absence of environmental turbulence, *L*_K_ and *L*_I_ are determined by the size of the trail source.

### B. Sensing range for chemosensing

Size limits for the functioning of chemosensing also apply to the sensing range. For example, in gradient climbing, the maximal distance up to which a chemical gradient remains uninterrupted is *L*_B_. Another factor affecting the range for gradient climbing is the diffusion time scale. For a typical compound to diffuse over *d* = 1 cm, it can take up to days (*t* = *d*^2^*D*^−1^ where *D∼* 10^−9^ m^2^s^−1^). This makes the signal irrelevant for many small organisms, because by that time they have moved elsewhere, been preyed upon, or have multiplied several times. Thus, gradient climbing is relevant only up to small distances. Similarly, for trail following, sensing range is limited to *L*_K_.

**FIG. 2.**
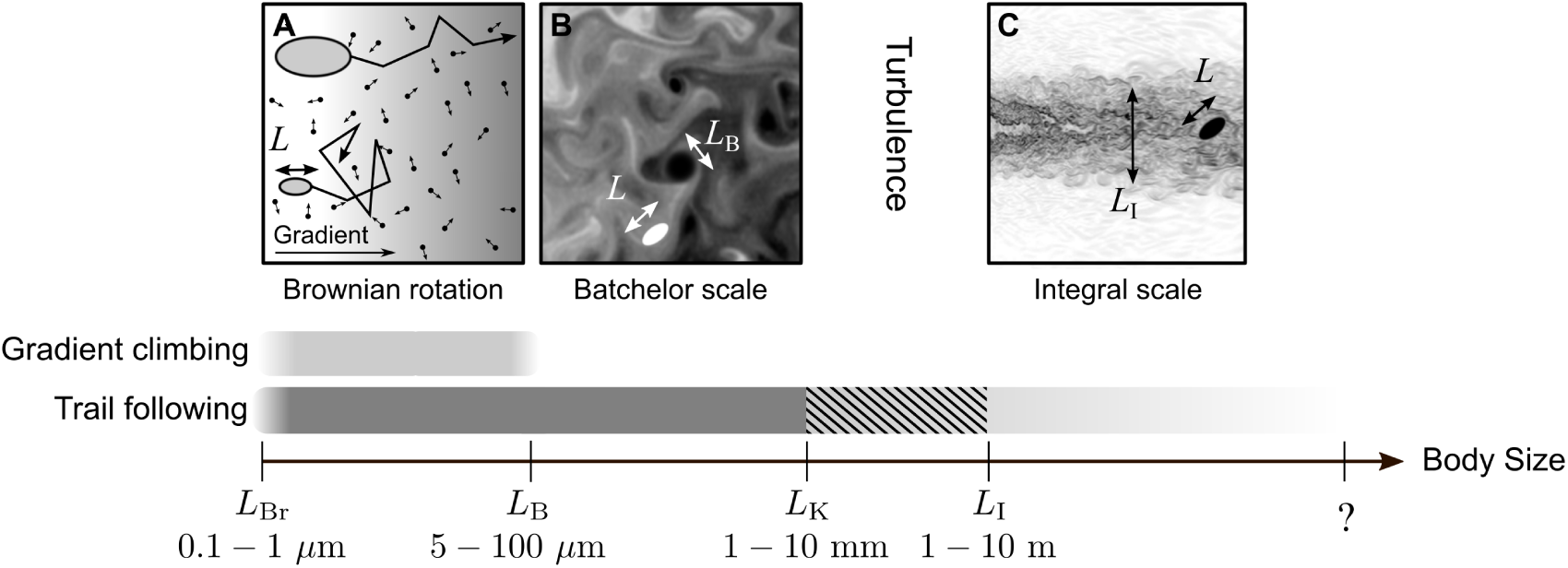
Body sizes over which chemosensing can be used effectively. A schematic illustration of Brownian rotation (A), Batchelor scale (B), and integral scale (C) is included at the top.

## IV. MECHANOSENSING

Any object moving in fluid generates a hydromechanical disturbance that can potentially be detected with the appropriate sensory apparatus [23]. For many small organisms such as zooplankton [23–25], it is the dominant sensory mechanism. Many fishes, especially in dimly lit environments, also rely heavily on mechanosensing using the lateral line organ [26]. The nature of a fluid disturbance generated by a target of size *L*_t_ swimming with a velocity *U*_t_ is largely determined by the dimensionless Reynolds number (Re), defined as Re = *L*_t_*U*_t_*/ν*, where *ν* is the kinematic viscosity [27]. For small Re, such as for most plankton, flow is dominated by viscosity and is laminar [28]. For large Re, such as for large fishes or mammals, inertia dominates, and the flow tends to be turbulent [29].

### A. Propagation of fluid disturbances

For a target passively sinking at low Re in unbounded fluid (e.g., the pelagic zone), the velocity (*u*) induced in the fluid decays with distance *r* as *u∼ r*^−1^ [23]. For a self-propelled target, the induced velocity decays as *u∼r*^−2^ [23]. Recent studies have shown that for breast-stroke swimming plankton and impulsively jumping copepods, *u* decays more rapidly as *u∼r*^−3^ and *u∼r*^−4^, respectively [30, 31]. At high Re, the fluid disturbance generated by a target becomes turbulent, if *L*_t_ is much larger than *L*_K_, resulting in a turbulent wake.

### B. Detection

Setae on the antennae of a copepod are classic examples of mechanosensors (Supplementary Figure 2). Setae sense velocity difference across their length, and activate when it exceeds a certain threshold *s* [25], defining setae sensitivity [32], typically between 10 and 100 *μ*m/s [23]. In unicellular organisms such as ciliates and dinoflagellates, a response occurs above a critical fluid deformation rate [24, 33], equivalent to a threshold velocity difference across the cell. In the lateral lines of fish, the working sensor is a seta-like kinocilium [34]. In general, mechanosensing requires a velocity differential on the organism’s body, as a result of fluid deformation. Given a sensitivity *s* of a mechanosensor of length *b*, embedded in fluid with deformation rate Δ (measured in s^−1^), the criterion for detection can be written as

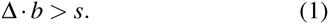

### C. Sensing range for mechanosensing

We estimate the sensing range *R* for the most relevant case of a self-propelled target. For *R* >> *b*, Visser [23] has shown that 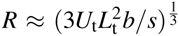. The swimming velocity of the target is related to its size by the empirical relation 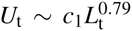 with *c*_1_ =6.5 m^0.21^*/s* [1]. For prey detection (*p* = 0.1), assuming that the sensor is about a tenth of the body size (*b* = *L/*10), we get

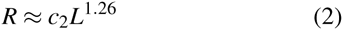

where *c*_2_ = 3.98 m^−0.26^.

From this estimate, a copepod of *L ∼*2 mm has a prey sensing range of about 1.5 mm. The exact scaling coefficient is determined by the organism’s morphology and the swimming characteristics of the target, but equation (2) provides a rough estimate. Like in chemical trail following, an upper limit of mechanosensing range *R* is set by the Kolmogorov scale, *L*_K_, above which turbulence disrupts the signal.

### D. Size limits for mechanosensing

The lower size limit for mechanosensing in the pelagic zone is dictated by inequality (1). We consider the case of a small prey individual detecting a larger predator (*p* = 10). For a target (predator) swimming with a velocity *U*_t_, fluid deformation scales as Δ *∼U*_t_*/L*_t_. Using again the empirical scaling of 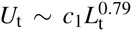 [1], and further using *L* = *L*_t_/10, we can deduce that

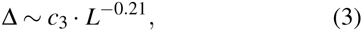

where *c*_3_ = 3.98 m^021^s^−1^.

To close the problem, we again use *b* = *L/*10. Combining (1) and (3), substituting *b* and using an intermediate value for *s* = 50 *μ*m/s, we get a lower size limit of *L >* 11 *μ*m. Thus we expect the lower size limit for an organism to use mechanosensing in the pelagic zone to be of the order of a few micrometers. Given the sensitivity of mechanosensing apparatuses, smaller organisms are unable to detect the hydromechanical disturbances relevant to their size.

The upper size limit of mechanosensing is prescribed by the same constraints as those for chemical trail following. The generated flows are disintegrated by turbulence at *L > L*_K_, rendering mechanosensing progressively less effective above organism sizes of around 1 cm. We also conjecture that like trail following, mechanosensing abilities may improve for organisms larger than the integral length scale *L*_I_.

## V. VISION

Simple functions of vision include differentiating light from dark, entrainment to a circadian rhythm [35], and orientation [36], while more complex functions involve navigation, pattern recognition, and food acquisition. Prey and predator detection from some distance requires sufficient image resolution. In general only two fundamental principles are used to build an eye: i) compound eyes, which comprise of a number of individual lenses and photo-receptors laid out on a convex hemispherical surface, ii) camera eyes with one concave photoreceptive surface where an image is projected through an optical unit (pinhole or lens).

### A. Light propagation in the marine environment

Given that a target is lit and visible, the reflected light must travel through seawater to reach the receiving organism. The intensity of light attenuates geometrically with distance *r* as *r*^−2^, and more steeply due to the added effects of scattering and absorption by solutes and seston [37]. In general, light intensity along a given path decreases as *e*^−α^*^r^* where α (measured in m^−1^) is called the absorption coefficient [38].

### B. Physiological limits to eye size

The resolution of the compound eye is limited by the size of ommatidia (photoreceptor units in compound eyes). They cannot be reduced in size to achieve a resolution better than 1*^°^*[39]. Thus, camera eyes, which we consider in the following, outperform compound eyes in compactness [39, 40]. The functioning of a small eye is limited by two constraints. First, a smaller eye captures less light. Second, a smaller eye has lower resolution: the photoreceptive units constitute the smallest components in an eye and are based on opsin molecules, the universally represented light-capturing design in the animal world [41]. Thus, the width of a photoreceptor *d*_*p*_*≈* 1 *μ*m [42] is an absolute limiting factor for any eye design. Therefore, *n* pixels amount to a retina diameter of *d ≈ n*^1/2^*d*_*p*_. Considering a minimal required resolution for a usable image-forming eye to be 100^2^ pixels, the corresponding retina would have a diameter *d ≈* 0.1 mm. Depending on the eye-to-body size ratio, this corresponds to an organism of around *L ≈* 1 to 3 mm.

Arguments for an upper size limit for eyes are not evident on physical grounds. The largest known marine animals carry eyes (see Discussion). However, the higher resolution and sensitivity resulting from larger eyes do not necessarily yield a larger sensing range as it may be limited by turbidity, as we discuss next.

### C. Visual range

The visual range of an organism can be estimated by considering the properties of a (pin-hole) camera eye, following an argument by Dunbrack and Ware [43]. We use Weber contrast *C* = (*I−I_b_*)*/I_b_*, where *I* and *I*_*b*_ are the intensities of the target and the background, respectively. The maximal distance *R* at which a predator can discern a prey individual of size *L*_t_ requires that the apparent contrast *C*_a_ of the target matches the contrast threshold of the eye, *C*_th_. The inherent contrast of the target, *C*_0_ declines with distance *r*, yielding [38]

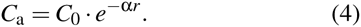

*C*_th_ is a declining function of the number of visual elements *n* involved in perceiving the target:

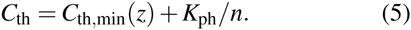

This formula is partly based on Ricco’s law [44] that expresses the inverse proportionality between *C*_th_ and *n*, and is supplemented by adding the minimum contrast threshold *C*_th,min_ to represent saturation of the contrast at a minimal value [45]. *C*_th,min_ varies in different environments and, in particular, depends on the available backlight at a given depth *z*.

The number of visual elements *n* involved in image detection is equal to their density, σ (measured in m^−2^), times the projected image area. Assuming *R* is large relative to the eye ball diameter *L*_eye_, we can deduce 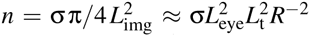(Supplementary Figure 3). Noting the universal size of the opsin molecule across species, we may assume that σ is independent of eye size. Introducing the ratio *a* = *L*_eye_*/L* [46] and using *p* = *L*_t_*/L*, we get *n* = σ*a*^2^ *p*^2^*L*^4^*R*^−2^. The range *R* is determined by the condition *C*_*a*_ *≥ C*_th_:

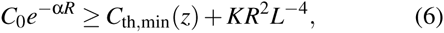

where *K* = *K*_ph_σ^−1^a^−2^ *p*^−2^ is a constant characterizing the photoreceptor sensitivity, *K*_ph_/σ, eye-to-body-size ratio, *a*, and size preference, *p*. Sample solutions for the condition *C*_*a*_ = *C*_th_ yield the range *R* at a given body size *L* (Figure 3A). Isolating *R* from Eq. (6) is impossible; however, asymptotic solutions can be derived for two limits:

(i) “Clear-water limit”: when α → 0, *R* is limited by the eye’s resolution; thus, *R∼*[(*C*_0_ *-C*_th,min_)*K*]^1/2^*L*^2^.
(ii) “Turbid-water limit”: when *C*_0_ >> *C*_th,min_ *KR*^2^*L*^−4^; thus, *R∼*(ln*C*_0_*-*ln*C*_th,min_)/α. *R* is independent of *L* and only limited by the sensitivity of a visual element, *C*_th,min_.

Generally, the visual range decreases if light is reduced, e.g., at large depth *z*, leading to a higher *C*_th,min_ [cases (i),(ii)]; or if the turbidity is strong (larger α) [case (ii)]. The cross-over between the two limits occurs when *L∼ L_x_ ∼*α^−1/2^ (Supplementary text). The visibility range in pure water for light of 550 nm is theoretically estimated at 74 m [47], and measurements in the open sea range from 44-80 m [48]. The visual range has also been predicted in more elaborate models [49].

## VI. HEARING

Sound propagates through the ocean as pressure waves, resulting in alternating compression and rarefaction of water in regions of high and low pressure, respectively. Any form of hearing must detect sound waves by converting them into vibrations of an organ that stimulates nerve cells. In fishes, sound waves displace sensory hairs against the calcareous *otolith*, and this relative motion is detected. By contrast, in mammalian ears, sound waves excite the tympanic membrane (ear-drum), the motion of which is sensed by ciliary hairs in the cochlea.

Most sounds relevant to ocean life, except echolocation, fall into the range of a few hertz up to a few kilohertz. Sounds generated by marine animals due to rapid movements or for communication, have frequencies rarely exceeding 1 kHz [50]. Communication by marine mammals usually consists of a burst of clicks or of whistles (4-12 kHz), while the echolocating signals of odontoceti range between 20 and 200 kHz [51].

### A Underwater sound propagation

As sound waves travel through a medium, sound intensity attenuates with distance from the target *r*, due to two processes: (i) geometric spreading (*r*^−2^ in open space), and (ii) absorption in water. The latter is frequency dependent: 1 dB/km at 10 kHz, but only 10^−4^ dB/km at 100 Hz in seawater^1^ [38]. Sound is therefore only weakly attenuated in seawater, and it can potentially carry information over large distances.

### B. Lower limit for sound detection

Detection of sound requires either an organ of significantly different density than that of water (e.g., the otolith), or a large detector array (e.g., auricle and drum), to allow detection by responding to spatial gradients of particle displacement [38]. A density contrast organ such as the otolith has to move relative to the surrounding fluid, as explained above. Motions in small sound-sensing organs (operating at low *Re*) are inherently more damped by viscosity than larger ones, impairing the practicality of sound detection by small organisms. Without high density contrast in the hearing organ, the detector array and thus the organism would have to be at least as long as the wavelength of sound (15 cm at 10 kHz). Thus hearing – with or without a density contrast organ – is impractical for pelagic organisms smaller than a few centimetres.

Many fishes have swim bladders (sometimes connected to the otolith-containing cavity through bony connections called the *Weberian ossicles*) that transduce pressure waves to mechanical motion and act as displacement amplifiers for sound via resonance [38, 52]. Similarly, odontocetes use the fat-filled bones of their lower jaw as an amplifying cavity [51]. Swim bladders are air-filled structures that amplify sound maximally when in natural resonance with the sound waves [38]. Frequencies very different from the resonance frequency of the swim bladder do not amplify well, and may even be damped if too different [38]. Based on an assumption of a spherical, air-filled swim bladder, the resonance frequency, *f*, can be approximated [38] as

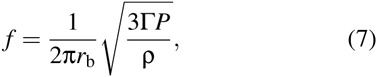

where *P* is the depth-dependent hydrostatic pressure, *r*_b_ the radius of the swim bladder, ρ the density of sea water, and Γ the adiabatic exponent (*∼*1.4 for air); *r*_b_ is typically around 5-10 % [53] of the body size *L* of the fish. Using *r*_b_ = *L/*10 for a conservative estimate, *L* would need to be at least 3 cm at the sea surface, in order to amplify the high-frequency end (1 kHz) of the ambient underwater sound spectrum, and *L* = 11 cm at a depth of 100 m (Figure 3B). To hear the more typical lower frequencies, *L* would have to be larger still. Thus, we approximate that the lower body size limit for detection of sound using swim bladders is around a few centimetres.

**FIG. 3.**
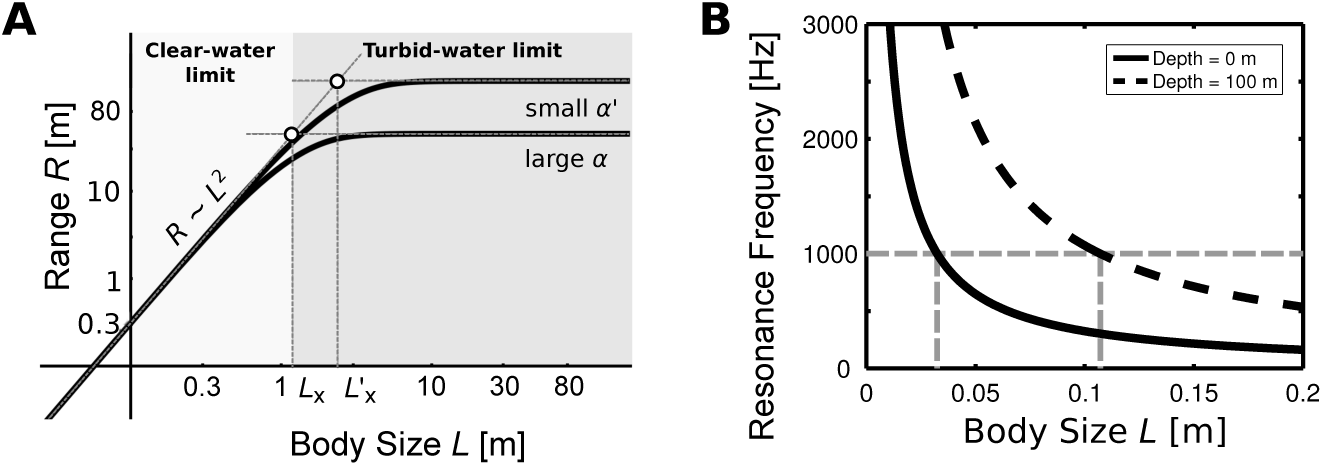
**A**: Visual sensing range scales with body size, *L*, as *R ∼ L*^2^ in the clear-water limit (*L << L_x_*) and as *R ∼* constant in the turbid-water limit (*L >> L_x_*). Parameters are *C*_0_ = 0.3,*C*_th,min_ = 0.05 (adopted from [43]), *K* = 2.5 *×* 10^−4^ m^2^, α’ = 0.04 m^−1^ [69] (and α’ = 0.01 m^−1^ for comparison). **B**: Relationship of body size and resonance frequency based on Equation (7) and using a swim bladder size *r*b = *L/*10 for an individual at the surface (solid curve) and at 100 m depth (dashed curve). The dashed (grey) horizontal line indicates 1 kHz, below which most sounds generated by marine life are found.

## VII. ECHOLOCATION

Echolocation is an active sensing mode, in which the organism emits clicks in the ultrasonic range and interprets the environment based on the echoes of these clicks. Echolocation is common in odontocetes (toothed whales) and is generally used for orientation and prey detection. The generation of echolocating signals in toothed whales is associated with the nasal passage leading up to the blowhole and takes place in the phonic lips. Taking into account the anatomical structures, the dominant frequency can be estimated as the resonance frequency of a Helmholtz oscillator [54]. The diffraction limit sets a resolution limit to λ/2π, where λ is the characteristic wavelength of the click [38]. Odontocetes produce clicks with peak energies at frequencies in the range of 20 to 200 kHz [51], the resulting resolution lies between 1 to 8 mm. Using an intermediate value (5 mm), and assuming that the target is at least one order of magnitude larger than the smallest resolvable feature, we get a minimal target size of 50 mm. Echolocation is typically used for prey detection, so *p* = 0.1. Thus we get a lower body size limit for an echolocating organism to be *L≈* 500 mm. It also implies that objects smaller than about 1 mm do not scatter sound signals in the frequency range we are considering, allowing echolocation to be useful in turbid waters where vision is severely restricted.

### A. Sensing range

The generated acoustic signal first travels through water, is then partially reflected by the target, and the remainder of the signal (minus attenuation) travels back to the organism. Emitted sound intensity, *I*_*e*_, is thus reduced by the processes of reflection and geometric divergence, causing signal intensity to attenuate as (2*r*)^−2^*e*^−2^*μr*. The strength of the returned signal must exceed the threshold intensity for detection in the ear, *I*_*r*_ = *I*_0_. Assuming that ear threshold sensitivity is independent of *L*, but that emitted sound intensity *I*_*e*_ and carrier frequency scale with *L*, the sensing range can be estimated as (Supplementary Text for details)

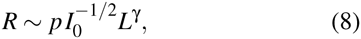

where *p* = *L*_t_*/L* is the size preference ratio and the exponent Γ lies between 2.125 to 2.5 that compares reasonably well with data. The scaling factor can be estimated from data describing the echolocation range of small marine mammals (Supplementary Text).

## DISCUSSION

We have attempted to synthesize an understanding of how physiology and the physical environment enable and constrain an aquatic organism’s ability to gather information from its surroundings. By reducing the relevant physical mechanisms to their simplest forms, we have identified the most pressing constraints on the functioning of various senses. Our goal has been to explain the transition from one dominant sense to another with changing body size, as observed in nature. A comparison of the predicted size limits with those observed in nature supports our analysis (Table I, Figure 4). The predicted size ranges correspond well with known minimal and maximal sizes of animals using a specific sense. Size limits of a sense do not imply that an organism cannot detect the signal outside the limits at all, but rather that beyond these limits, the usefulness of the sense is compromised in comparison with other senses.

We could not conceive any upper size limits on physical grounds for chemosensing, mechanosensing, hearing, and vision. Indeed, the largest known organism in the ocean, the blue whale (*L* = 30 m), is known to use all of these senses. Chemosensing is the only sense available to the smallest organisms, and its theoretical lower size limit (*L*_Br_*∼*10^−7^*−*10^−6^ m) is consistent with the smallest known motile organisms (bacteria, *L* = 0.8 *μ*m [16]). Chemosensing is presumably slightly impaired due to turbulence in intermediate size ranges, in which integration of multiple senses such as mechanosensing and vision might be very useful. Chemosensing for trail following is an important sensory mode for large bony fishes [61] and sharks [62], which have sizes larger than *L*_I_.

**TABLE I.**
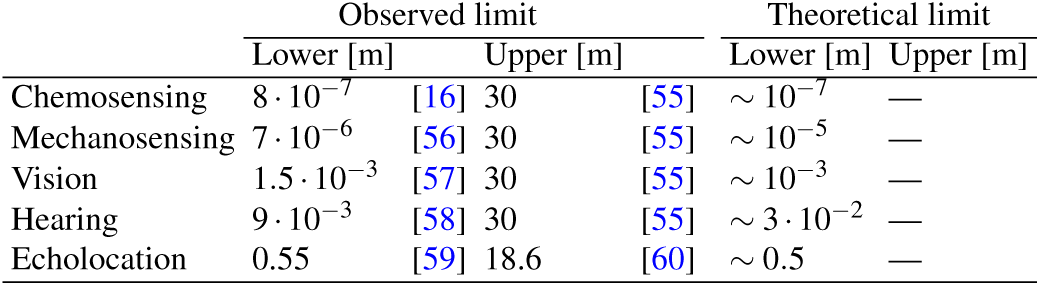
Lower and upper size (body length) limits for various senses. Predicted theoretical limits denote orders of magnitude.

**FIG. 4.**
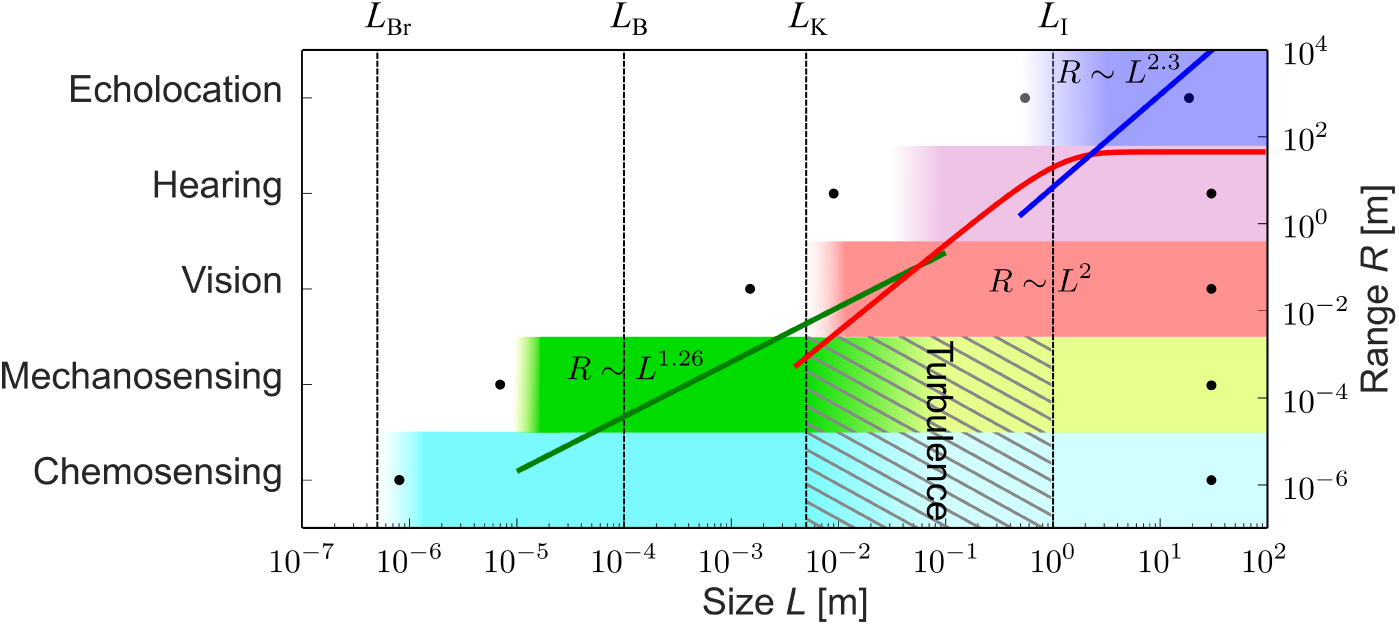
Upper and lower body size limits and ranges for different senses. Dots denote the largest and smallest sizes known to employ a given sense, and shaded rectangles show the theoretical estimates of the size range in which a sense is expected to work. Green, red, and blue curves show the theoretical scaling of sensing range with size for mechanosensing, vision, and echolocation, respectively.

The theoretical lower limit for mechanosensing in the pelagic environment is a few micrometers, in the realm of protists; to our knowledge, marine protists sized 7-10 *μ*m are the smallest pelagic organisms known to use mechanosensing [56]. However, it is only the lower limit for pelagic zones; smaller bacteria are known to be able to sense mechanical stresses when getting in contact with a solid body [63]. Large copepods and small fish occupy the size range where mechanosensing starts becoming less effective. Its use by fish is demonstrated in many species using lateral lines to find prey and sense flows [26]. Larger fish receive a poorer signal quality due to turbulence, and for this reason some larger sharks are known not to use lateral lines for prey detection [64]. Some marine mammals (seals and sea lions) have the ability to follow turbulent trails using their mystacial vibrissae [65], likely due to being larger than the integral length scale set by the target.

The camera eye takes records for both the smallest and the largest eye: the smallest image forming eyes (and body sizes) are found in the fish *Schindleria brevipinguis* (*L ≈*7 mm [66]), and the pygmy squids (*L ≈*1.5 mm [57]), which compares well with our predicted size limit^2^. The largest known eye belongs to the giant squid, featuring eye-balls up to 30 cm in diameter [67]. Eyes are also found in the largest known species (whales), implying that there is no upper body size limit for image-forming vision in marine animals. For hearing, the theoretical lower body size limit is found to be a few centimetres. Some fishes are able to manipulate the resonance frequency of swim bladders by changing their membrane elasticities [68]. By hearing outside the resonance frequency, fish larvae of a few millimetres (*L≈* 9 mm) have been shown to react to sounds [58]. Note that these fishes inhabit shallower waters, where hearing is feasible at smaller sizes (Figure 3B). For echolocation, the predicted lower limit (*∼*0.5m) is close to the observed smallest size among echolocating marine mammals (Commerson’s dolphin, [59]).

Upper limits of sensing ranges are dictated by degradation of signal-to-noise ratios via absorption, geometric spreading (divergence), or environmental disturbances. For chemical gradient climbing and mechanosensing, the signals are randomized beyond a characteristic distance given by *L*_B_ and *L*_K_, respectively. For mechanosensing the range scales as *R∼ L*^1.26^ (Figure 4). When mechanosensing can no longer extend its range, vision becomes a viable solution. Visual sensing range in clear water scales as *R∼L*^2^, but cannot exceed the limit set by turbidity. Even in clear waters, vision cannot exceed the range of roughly 80 m. Here, vision may be complemented by hearing and echolocation mainly because sound is capable of travelling large distances in sea-water without significant attenuation. Although we could not develop a scaling for hearing range, we could determine the sensing range of echolocation, which scales approximately as *R∼ L*^2.3^ and is as large as kilometres for larger organisms, comparing well with the known range of marine mammals.

The question arises whether there is a general pattern underlying the size structure of primary sensory modes. For instance, can the transitions between senses be related to metabolic demand? Kleiber’s law requires that an organism consumes energy at a rate proportional to *L*^9/4^[3]. This demand must be fulfilled by maintaining a sufficient clearance rate [4], a function of the swimming velocity *V ∼L^x^* and sensing range *R ∼L^y^* with positive exponents *x, y*. Thus, the clearance rate also increases with *L*. The exponent *y* appears to increase going up the senses axis (Figure 4). With increasing size and metabolic expenditure, an evolutionary pressure arises to extend the sensing range by investing into a more effective sensory strategy, causing the transition from one to the other primary sensing mode. However, rather than being governed by cost efficiency, it seems more plausible that the transitions between senses are set by the physical limitations of signal generation, transmission and reception. To exemplify, carrying larger eyes can improve resolution and thus extend the sensing range, but beyond a critical (eye) size, increased performance is rendered ineffective due to the clear water limit of the visual range. So a transition is necessitated by the required increase in sensing range, achieved by echolocation.

We have combined biological knowledge, physiology and physics to describe the abilities of the sensory modes in ocean life, from bacteria to whales. Our treatise demonstrates how body size determines available sensing modes, and thereby acts as a major structuring factor of aquatic life. When interpreting the scalings and limits we propose, note that our purpose is to provide first-order approximations based on first principles. Further research is needed to evaluate each of the senses in more detail and to gather more data to examine the arguments presented here. We hope that this work may serve as a starting point for future explorations on sensory modalities and their hierarchical structures.

1 The decibel level is defined via *I*_dB_ = 10 log10 (*I/I*_0_), where *I* is the sound intensity and *I*_0_ is a reference frequency.

2 The smallest compound eyes are found in the genus *Daphnia*, but their image quality is questionable, see Supplementary Text.

## AUTHOR CONTRIBUTIONS

EAM and NW contributed equally to the study, collected data, developed models, and wrote the manuscript. NSJ, CL, KHA and AV collected data and helped drafting the manuscript. All authors participated in the design of the study and gave final approval for publication.

## ACKNOWLEDGMENTS

We thank Hanna Rademaker, Julia Dölger and Anders Andersen for helpful discussions, and Thomas Kiørboe for comments on the manuscript. We thank anonymous reviewers for helpful comments that improved the manuscript. The Centre for Ocean Life is a VKR center of excellence supported by the Villum foundation (EAM, NW, CL, NSJ, KHA, AV). The work is part of the Dynamical Systems Interdisciplinary Network, University of Copenhagen (EAM). Partial financial support for CL was provided by EURO-BASIN (FP7, Ref. 264933).

## Supplementary Information: Size Structures Sensory Hierarchy in Ocean Life

**FIG. 1.**
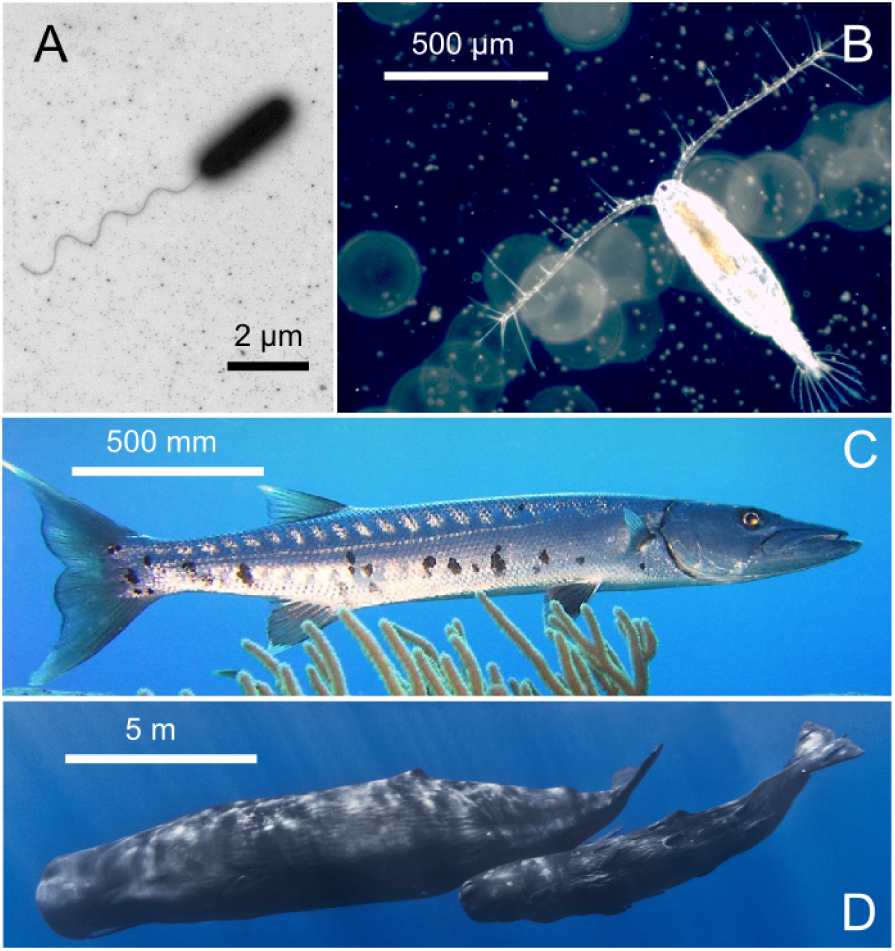
Dominant sensing modes. change with increasing size of the organism: (**A**) Small organisms like bacteria (e.g., *Vibrio alginolyticus*) use chemosensing, and move up or down the gradients of chemicals (image courtesy Kwangmin Son and Roman Stocker, MIT). (**B**) Millimetre sized organisms like copepods (e.g. *Acartia tonsa*) use hydromechanical signals to detect predators and prey in the vicinity (image courtesy of Thomas Kiørboe, DTU). (**C**) Larger organisms like fish (e.g. great barracuda *Sphyraena barracuda*) are often visual predators. (**D**) Toothed whales (e.g. *Physeter macrocephalus*) use echolocation. Images in panels C,D are in public domain.

### I. CHEMOSENSING

#### A. A note on chemical contrast

An absolute upper limit on sensing range is dictated by the requirement of sufficient chemical contrast. Chemosensing requires spatial variations in signal strength that can be detected and gradients therein tracked. However, chemical gradients tend to become eroded with time to background level. The upper limit chemosensing range is not only related to the size and sensory ability of the organism, but also to the nature of the chemical substrate and its degradation in the environment due to microbial action or chemical reactions. Thus, while it is clear that an upper limit to chemosensing range exists, it is not possible to quantify it.

**FIG. 2.**
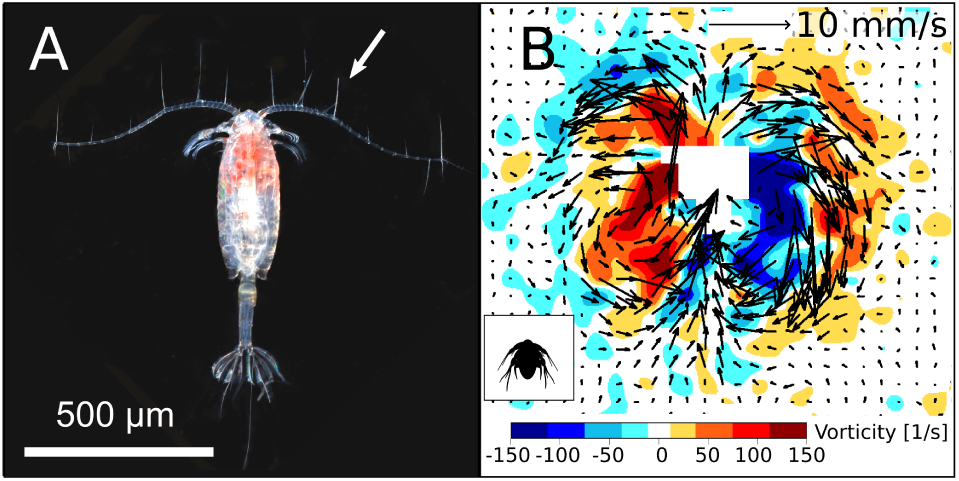
Mechanosensing. ***A:** Dorsal view of an adult Acartia tonsa*, showing the antennules covered with mechanosensory setae, one of which is marked with an arrow (image courtesy of Erik Selander). **B:** Flow disturbance created by a swimming *Acartia tonsa* nauplius, visualized in the form of velocity vectors and vorticity contours.

## II. VISION

### A. Size limit for compound eyes

The compound eye is hemi-spherical in shape and subdivided into light-detecting units called *ommatidia*. Ommatidia are conical in shape and cover the surface with an opening of width *δ*. Given that the eye has a radius *r*, the visual acuity of an ommatidium is given by

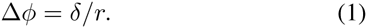

The number of ommatidia covering the hemispherical eye surface may be estimated as the ratio of the eye surface, around 2*πr*^2^, and the surface element covered by an ommatidium, around *r*^2^Δ*φ*^2^,

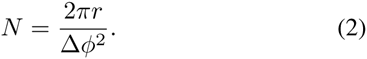

Increasing the number of ommatidia, *N*, enlarges the image-resolution of the eye; however, as *δ* is decreases, diffraction effects becomes increasingly important. Thus, minimization of ommatidia in compound eyes is limited due to diffraction limits, see [1, 2]. Considering this trade-off, the optimal width of the ommatidia can be estimated [1], yielding

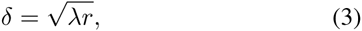

where *λ* = 400nm is the wave length of blue light.

Substituting Eqs. (1) and (3) into Eq. (2), we obtain the resolution of an eye with optimal ommatidia, we have

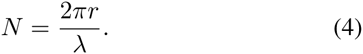

The size of an optimal compound eye is then

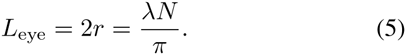

The size of the optimal compound eye with a reasonably useful resolution of *N* = 100^2^ pixels should be *L*_eye_ = 1.2 mm, corresponding to roughly *L∼* 1 to 3 cm (depending on the size ratio of eye to body). By comparison, some of the smallest organisms carrying compound eyes are *Daphnia*, with adults ranging from 1 to 5 mm [3]. Optimality of the eye, Eq. (5), then implies a resolution of *N∼* 10^2^ pixels – however, this is a resolution which barely produces a usable image.

### B. Sensing range

The sensing range condition in the main text is given by

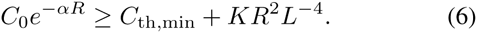

Rescaling the sensing range, 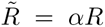 and the size, 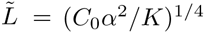 where *C*:= *C*_th,min_*/C*_0_, this becomes

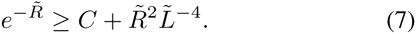

The clear-water limit corresponds to small 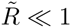 « 1, yielding

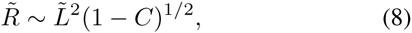

and the turbid-water corresponds to large 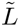, yielding

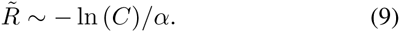

These expressions match the ones presented in the main text.

Letting the two expressions for the rescaled sensing ranges (8) and (9) be similar, we arrive at the condition for the crossover between the two regimes:

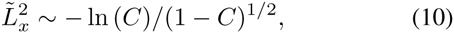

which in the original unscaled variables becomes 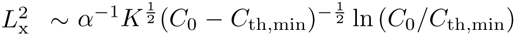 or, to leading order,

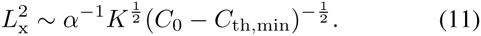

The clear-water limit occurs for *L << L_x_* and the turbid water limit for *L >> L_x_*. Thus, the turbid limit is reached in the limit of large *α*, large (*C*_0_ *-C*_th,min_), or small sensitivity *K*, respectively.

Another (rough) estimate of the minimal body size, for which vision is still marginally meaningful, might be feasible from the condition that *L∼R*. This condition has at most two solutions, whereas the minimal solution is *L ≈*[*K/*(*C*_0_*-C*_th,min_)]^1/2^. A precise determination of this estimate of the smallest animal carrying an eye is, however, difficult due to the unknown scaling coefficient in this estimate and uncertainties concerning parameter values.

**FIG. 3.**
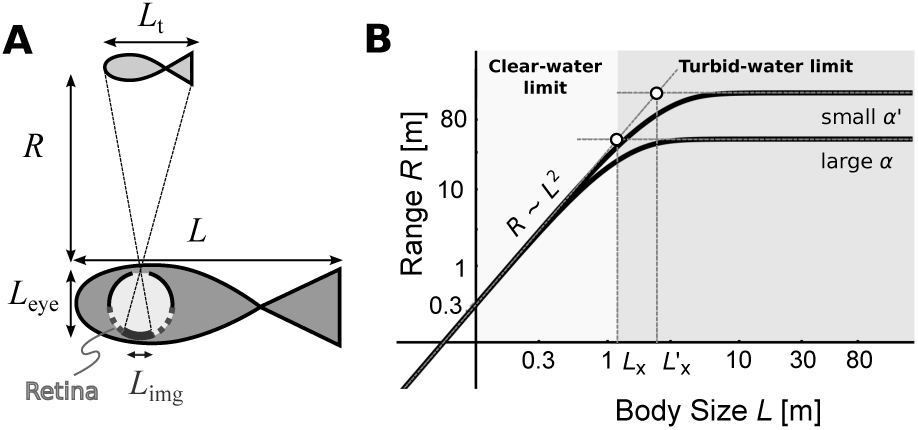
Vision. A: An organism of body size *L*, with an eye of size *L*_eye_, detects a target of size *Lt* at a distance *R* if the apparent contrast of the target is equal or larger than the threshold contrast of the organism’s eye. **B**: Maximal visual sensing range scales with body size *L*: like *R ∼L*^2^ in the clear-water limit (*L «Lx*) and like *R ∼*constant in the turbid-water limit (*L »Lx*). Parameters are *C*_0_ = 0.3, *C*_th,min_ = 0.05 (adopted from [4]), *K* = 2.5 *×* 10^−4^ m^2^, *α* = 0.04 m^−1^ [5] (and *α*’ = 0.01 m^−1^ for comparison).

## III. ECHOLOCATION

### A. Scaling argument for sensing range

We estimate how the range of echolocation scales with body size *L* based on three assumptions: i) the threshold sensitivity of the ear *I*_0_ is independent of organism size *L* [6], ii) the emitted sound intensity I_e_ scales with size: *I*_*e*_ *∝L*^3φ^ where 3/4 *< φ <* 1, and iii) the carrier frequency of the signal depends on *L* (see [7]).

The generated acoustic signal first travels through water, is then partially reflected by the target, and the remainder of the signal travels back to the organism. *I*_*e*_, is thus reduced by two processes:

i) ***Reflection.*** The signal is reduced upon reflection from the target and the reflected intensity is proportional to the target area which scales as 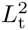.

ii) ***Attenuation.*** Sound intensity decreases with distance as *r*^−2^ due to geometric divergence. It is further attenuated exponentially due to absorption in the seawater.

Together, the signal intensity attenuates as (2*r*)^−2^*e*^−2^*μ^r^*, where the factor 2 is due to the doubled travel distance. Geometric attenuation strongly dominates over the absorption processes, thus, 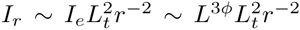. The strength of the returned signal must exceed the threshold intensity for detection in the ear, *I*_*r*_ = *I*_0_, yielding a sensing range 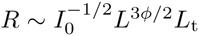. Introducing the size ratio *p* = *L*_t_*/L*, we arrive at

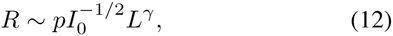

where the exponent *γ* = 1 + 3*φ/*2 lies between 2.125 to 2.5. The scaling factor depends on unknown parameters, but can be estimated from data describing the echolocation range of small marine mammals. The resulting scaling coefficient (including *p/I*_0_) is 6.47 m for *γ* = 2.5, and 9.79 m^−1.125^ for *γ* = 2.125.

Figure 5 compares the scaling for Eq. 12 with data available for dolphins [8–12]. There is considerable scatter in the data, yet we recognize that the prediction compares with the data reasonably well.

### 1. Signal attenuation

We detail our estimates for the effects of attenuation due to geometric divergence and absorption processes in sea water. First, we discuss the effect of absorption processes on the transmission of pulses. To begin, we note that the absorption coefficient *μ* is frequency dependent. Each pulse is transmitted and characterized by its center (or carrier) frequency *f*_*c*_, which is also the dominant frequency of the pulse spectrum. We may disregard all other frequencies and thus the dispersion of the transmitted pulse, leaving us with the task to find the absorption coefficient for *f*_*c*_. The attenuation of sound in seawater is a complex molecular process which occurs both due to viscous absorption generated by particle motion, but also due to molecular relaxation processes by Boric acid and Magnesium sulphate. A formula for the frequency dependent absorption has been devised [13]. However, this relation is too complicated for our purposes as we desire to establish a simple asymptotic scaling relation between *f*_*c*_ and *μ*; indeed, the data is well *parameterized* by 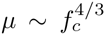(see Section 3 below and Figure 5). Further, it is known that *f*_*c*_ depends on body size; experimental data [7] for dolphins (excluding river dolphins), allows us to heuristically deduce a scaling dependence for the absorption, *f*_*c*_ ≈ 370 m^3/4^ s^−1^ × *L*^-3/4^ (see Section 2 below). Combining these two scalings, we obtain for the absorption coefficient (decibel/meter) *μ ≈*10^−2^*L*^−1^. Finally, since the fitted data is measured in the logarithmic decibel scale, the attenuation factor due to absorption converts to 10^-0.1^*μ*(*L*)*×*^2^*R*. Summed up, the intensity is reduced by a factor *I*_r_/I_e_ ∼ R^−2^10^-0.001^*×*^2^*R/L*. However, further analysis shows that the effect of damping is negligible when compared to the geometric divergence. Thus, the reflected sound intensity simplifies to 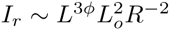.

### 2. Center frequency

Center frequencies of echolocation signals have been measured for dolphins [7], shown in Figure 4A. The two river dolphins discussed in [7] are excluded from our analysis, since dolphins in such environments operate at different frequencies to adapt for sound transmission in non-free environments. We fitted the relation between the body mass *w* and the center frequency by *f*_*c*_ *≈* (368.7 m^0.26^ s^−1^) *× m*^-0.26^. Since the mass scales as *w ∼ L*^3^, we obtain *f*_*c*_ *∼ L*^-3/4^.

**FIG. 4.**
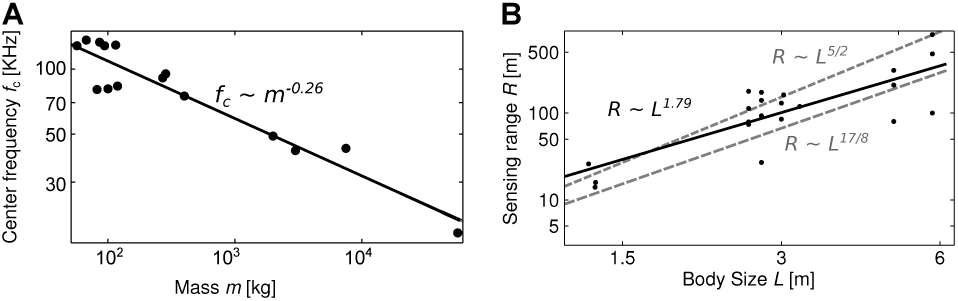
Echolocation. A: Power law fit for echolocation center frequencies of dolphins. Data from [7]: *fc* = (17.5, 42, 43, 75, 49, 95, 83.4, 80.4, 91, 81, 128, 136, 129, 133, 128) kHz; *m* = (57000, 3000,7500, 400, 2000, 285, 119, 82, 270, 100, 94, 67.5, 115, 86, 57) kg.**B:** Comparison of the predicted echolocation sensing range (dashed grey) with data (black dots), which scales like *R≈*14.2 m^-0.79^.*L*^1.79^(black line, least squares fit).

For comparison, note that the frequency with maximal intensity produced in the nasal sac is approximated by the Helmholtz frequency [14]:

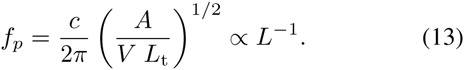

where *A*, *V*, *L*_t_ are the area, volume and length of the nasal sac. Given that *m* is proportional to *L*^3^, the scaling observed in Fig. 4 appears to deviate somewhat from this theoretical estimate. The deviation may be explained by shortcomings of the simple Helmholtz oscillator model.

### 2. Sound absorption in marine environments

The authors in [13] derive a simplified equation of the form

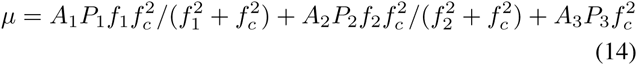

where the center frequency *f*_*c*_ is measured in Hz at the depth *z* in km. Further, they determine the following coefficients characteristic to the properties of seawater for boron and for magnesium,

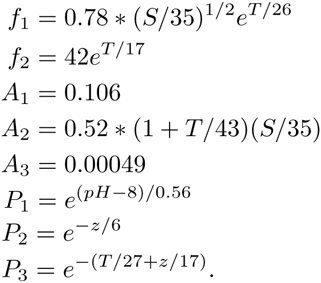

The scaling for the absorption coefficient *μ* is thus (decibel per meter)

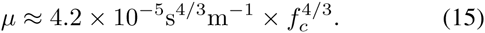

where the center frequency (s^−1^) is

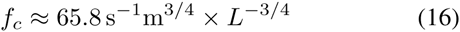

where we have used the relation mass w = *ρL*^3^ with *ρ* = 10^3^ kg m^-3^. Thus, we obtain (decibel per meter)

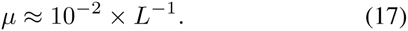

**FIG. 5.**
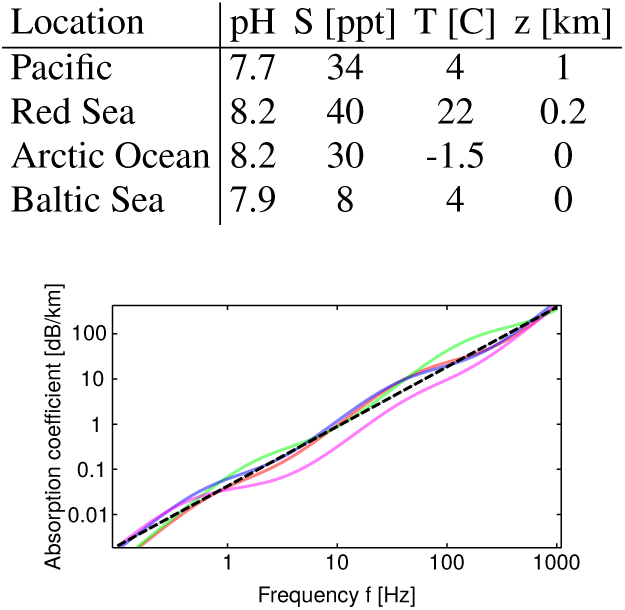
Power law fit for relation between frequency and sound absorption coefficient in the ocean. **Top:** Parameter values for *pH, S, T, z* for Eq. 14 valid for different ocean regions. **Bottom:** Absorption rates resulting from parameters for the various regions listed in the top table. Fitting the logarithmic data linearly (dashed line) over the frequency range of interest results in the asymptotic scaling relation *μ* [db/km] *≈* 0.0434 s^−4/3^km^−1^ *× f* ^4/3^[s^−1^].

## B. Assumptions underlying the scaling argument

The scaling argument for the range rests on assumptions supported by data only in part, which we review here for clarity:

(A1) the threshold sensitivity of the ear *I*_0_ is independent of target size *L*. This approximation is supported by audiograms (behavioral and auditory brain stem responses) of odontocetes [12, 15–18],

(A2) the emitted sound intensity that an animal produces scales with size: *I*_*e*_ *∝ L*^3^*φ* where 3/4 *< φ <* 1,

(A3) the carrier frequency of the sonar signal depends on size *L*.

Assumption (A3) seems fairly well corroborated, as already discussed in section A and B. Assumption (A2) states that the scaling exponent *φ* is allowed to vary in a small range corresponding to a sublinear volume dependence of the generating organ size which is a fairly reasonable assumption. Taking into account the considerable scatter of the data, we recognize that the prediction compares with the data reasonably well, as is evidenced in Figure 7 in the main text. However, better data is required to further underpin assumption (A1). Indeed, within the group of whales and dolphins we find no clear *size*dependence for the sensitivity threshold *I*_0_ [18]; but it would be desirable to obtain more data to solidify this assumption, as well as to identify a satisfactory physical or biological explanation for why the sensitivity is independent of body size, in contrast to other mammal groups [15–19].

